# Genetic Compensation Restores Embryonic Viability in Fatty Acid Synthase Mutants

**DOI:** 10.64898/2026.01.12.699145

**Authors:** Yooseong Wang, Milagros Rincon Paz, Xiao Fan, Harold E. Smith, Ricardo Roure, Anushka Patil, Crystal Shao, Christy Shao, Caitlyn Garcia, Ellie N. Brill, Ziqi (Kyra) Zheng, Sophia Monsalve, Samantha H. Bruner, Xiaofei Bai

## Abstract

Fatty acid synthase (FASN) is a key rate-limited, dimeric multi-enzyme complex in the *de novo* lipogenesis pathway. Each FASN monomer contains seven catalytic domains, which coordinate the stepwise conversion of acetyl-CoA into fatty acids. While FASN has been extensively studied in cultured cells, particularly for its oncogenic role, its functions in the germline and early embryonic development remain elusive. A major challenge is that the FASN dysfunction typically causes embryonic lethality in animal models, which complicates detailed functional analysis and the identification of compensatory genetic interactors during development. To overcome this limitation and identify novel genetic suppressors of the *FASN* gene, we utilized a temperature-sensitive allele, *fasn-1(g43ts)* (A1424T), in the genetically tractable model *Caenorhabditis elegans*, to conduct unbiased forward genetic screens. We isolated 22 suppressor lines that significantly restored embryonic viability in the *fasn-1(g43ts)* mutant at the non-permissive temperature. Using a combination of MIP-MAP genomic mapping and a customized bioinformatic pipeline, we identified six missense mutations in the *ptr-6* gene, which encodes a protein containing a patched domain associated with the Hedgehog signaling pathway. To validate this genetic suppression, we recreated one of the loss-of-function mutations, *ptr-6* (W701*), in the *fasn-1(g43ts)* background using CRISPR/Cas9 gene editing. Notably, *ptr-6(W701*)* robustly rescued the embryonic lethality and permeability defects caused by *fasn-1* loss-of-function. Taken together, our findings expand the genetic regulatory network of fatty acid synthase during early embryogenesis and highlight *ptr-6* and Hedgehog signaling pathway as potential genetic modifiers of FASN*-*associated developmental and metabolic disorders.

## Introduction

*De novo* lipogenesis (DNL) converts acetyl-CoA subunits from carbohydrate catabolism into long-chain fatty acids, which are essential for reproduction, embryonic development, and tissue morphogenesis ^1–4^. A central enzyme in this pathway, fatty acid synthase (FASN), functions as a homodimer, with each monomer containing seven enzymatic domains that orchestrate the sequential steps of fatty acid synthesis^5–7^. In vertebrates, loss of *FASN* results in embryonic lethality prior to implantation, while heterozygous *FASN* mutants arrest at multiple stages of early development^8^. FASN-dependent DNL is also required for tissue morphogenesis, including neuronal differentiation and the establishment of glial polarity in organoid models^2^. Although FASN has been extensively studied as an oncogenic biomarker and therapeutic target ^9, 10^, its roles in embryonic and tissue development remain elusive. The severe embryonic lethality and developmental arrest in vertebrate FASN-null models present major obstacles to addressing these questions^8, 11^, limiting *in vivo* functional analysis and preventing the identification of genetic modifiers that could rescue FASN-1-related defects.

*C. elegans* germline and embryos provide powerful models for studying FASN-dependent DNL during early development. During oogenesis and fertilization, somatic sheath cells surround the gonad and germline, providing essential nutritional support, mechanical forces, and signaling cues to ensure faithful fertilization and embryonic progression ^12–15^. Notably, the intestine, the primary nutrient-absorbing organ in *C. elegans*, communicates with sheath cells through channel proteins, such as gap junction complexes, enabling the direct transfer of small molecules, including lipids and fatty acids, to the germline during reproduction^12, 16–18^. In this study, we leveraged available genetic tools, including the temperature-sensitive mutation *fasn-1(g43ts)*, forward genetic screens, and an auxin-inducible degradation system (AID), to identify and validate novel genetic modifiers that suppress developmental defects in *fasn-1* mutants. We isolated 22 candidate suppressor lines from EMS-mediated forward genetic screens and identified six missense mutations in the hedgehog signaling pathway-related *ptr-6* gene. Additionally, we validated the loss-of-function effect of the *ptr-6* alleles during embryogenesis by regenerating the allele using CRISPR/Cas9 in the *fasn-1(g43ts)* background. Our current study suggests that the loss-of-function of the *ptr-6* gene could significantly increase permeability integrity, which may contribute to the impaired eggshell and embryonic defects observed in *fasn-1(g43ts)* mutants. The screening pipeline established here provides a framework for identifying genetic modifiers and signaling pathways that compensate for FASN-dependent defects, thereby advancing our understanding of how fatty acid synthesis and somatic tissue metabolism influence reproductive success and early development.

## Results

The CRISPR-edited *fasn-1(g43ts)* allele causes embryonic lethality and eggshell formation defects at the non-permissive temperature.

Our previous work identified a causative mutation in *fasn-1(g43ts)* as an Ala1425Thr missense mutation, which is a highly penetrant temperature-sensitive allele^19, 20^. Unexpectedly, the original *fasn-1(g43ts)* population displayed a high incidence of males, a phenotype absent in other *fasn-1* mutants, suggesting the presence of additional background mutations^19^. To obtain a clean *fasn-1(g43ts)* allele suitable for forward genetic screening, we regenerated the *fasn-1(g43ts)* allele in the wild-type background using CRISPR/Cas9 genome editing (Fig.1A).

**Figure 1.**
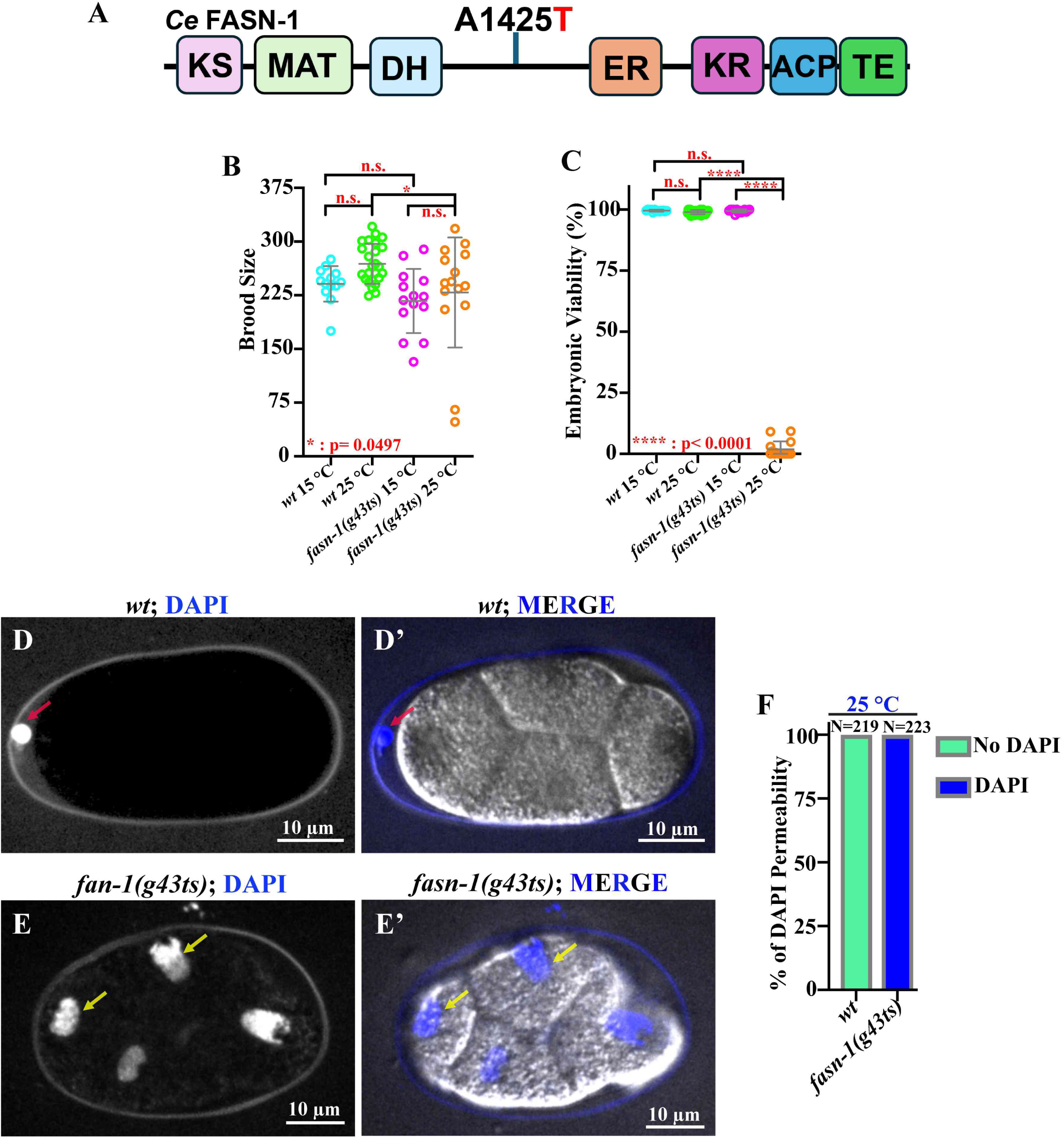
The CRISPR/Cas9-edited *fasn-1(g43ts)* allele is temperature-sensitive and compromises embryonic viability and eggshell integrity at the restrictive temperature. (A) Schematic diagram of *C. elegans* FASN-1 protein and the *fasn-1(g43ts)* (A1425T) allele. (B-C) Brood size and embryonic viability of the *fasn-1(g43ts)* allele at both 15 °C and 25 °C. (D-G) Representative images of DAPI staining of embryos to assess eggshell permeability. The red arrow in D-D’ indicates the stained polar body located outside the permeability barrier. The yellow arrows in E-E’ indicate the presence of stained zygotic chromatin. (F) Quantification of the DAPI staining of embryos at 25 °C. Scale bars are indicated in each panel.

The CRISPR-edited *fasn-1(g43sts)* allele behaved as a superficial wildtype at the permissive temperature (15°C) and did not exhibit secondary phenotypes such as elevated male frequency. To verify its temperature sensitivity, we assessed brood size, embryonic viability, and eggshell integrity at both 15°C and 25°C (Fig. 1B-F). Consistent with the original strain, brood size in the CRISPR-edited *fasn-1(g43ts)* mutant was identical to wildtype at 15 °C but was significantly reduced at 25 °C (Fig.1B). Additionally, the CRISPR-edited *fasn-1(g43ts)* mutant exhibited nearly 100% embryonic lethality at 25 °C, while embryos were fully viable at 15 °C (Fig.1C). DAPI staining further revealed strong penetration into the cytosol and zygotic chromatin in all tested *fasn-1(g43ts)* embryos at 25 °C (yellow arrows in Fig.1E-E’, F), indicating defective eggshell integrity. In contrast, in wild-type embryos at 25 °C, DAPI staining was restricted to the first polar body, which is outside the intact permeability barrier (red arrow in Fig.1D-D’, F)^19–21^.

### *The fasn-1(g43ts)* mutation causes developmental defects, but not the cellular localization of FASN-1::GFP

To assess the *in vivo* expression pattern of *fasn-1* in *C. elegans*, we generated multiple fluorescent reporter strains using CRISPR/Cas9 gene editing to tag the endogenous loci at either N- or C-terminus of FASN-1 (Fig.2, Supplemental Fig.1A-C’’)^12^. We observed that FASN-1::GFP was broadly expressed in tissues associated with lipid storage and metabolism, including the intestine and hypodermis (Fig.2B-B’’), as well as pharynx and excretory duct (Fig.2G-G’), vulva (Fig.2I-I’), and germline and gonadal sheath cells (K-K’). Notably, strong expression was also observed in early embryos, particularly within epidermal precursor cells at different developmental stages, which plays a crucial role in supporting proper *C. elegans* embryonic differentiation and patterning (Fig.2C-F).

**Figure 2.**
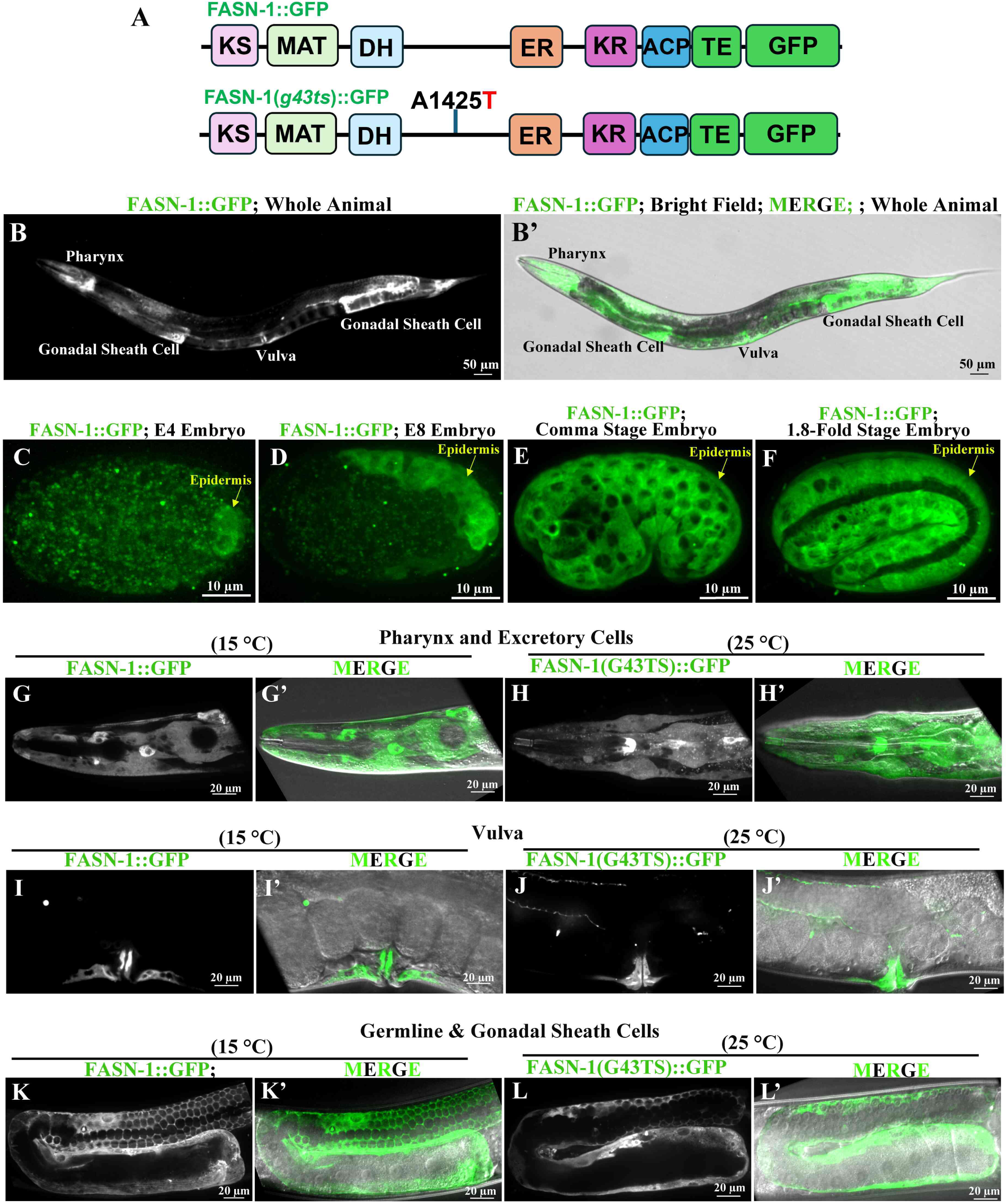
*fasn-1(g43ts)::gfp* does not disrupt the cellular localization of FASN-1::GFP. (A) Schematic diagram of endogenously tagged *C. elegans* FASN-1::GFP and FASN-1(g43ts)::GFP. (B-F, G-G’, I-I’, K-K’) FASN-1::GFP is expressed in a variety of cell types, including early embryos (C-F), pharynx (G-G’), vulva (I-I’), gonadal sheath cells (K-K’), and germline (K-K’). (H-L’) FASN-1(g43ts):GFP (green, H-H’, J-J’, L-L’) presents an identical expression pattern as FASN-1::GFP only control at the restrictive temperature, but with severe morphological defects in pharynx (H-H’) and vulva tissues (J-J’). DIC and GFP merged images are shown in panels G’-L’. Scale bars are indicated in each panel.

Additional fluorescent reporters, including GFP::FASN-1, co-localized with FASN-1::RFP across most cell types and tissues, suggesting that N- and C-terminus insertions did not impact the protein expression pattern (Supplemental Fig.1A-C’’). To determine whether *fasn-1(g43ts)* disrupts the temporal and spatial expression pattern of FASN-1, we generated the *fasn-1(g43ts)* allele by CRISPR/Cas9 genome editing into an endogenously tagged green-fluorescent reporter strain, *fasn-1::gfp*, which we designated *fasn-1(g43ts)::gfp* (Fig.2A). At the non-permissive temperature of 25 °C, we did not observe significant changes in the expression level or localization of FASN-1::GFP in most tissues, including pharynx (Fig.2H-H’), vulva (Fig.2J-J’), and germline (Fig.2L-L’). Taken together, the results from the fluorescent reporters suggest that the *fasn-1(g43ts)::gfp* mutation may compromise protein function and enzymatic activity, rather than altering the trafficking and cellular localization of FASN-1 *in vivo*.

### EMS-based forward genetic screen to identify suppressors of *fasn-1(g43ts)*

To identify genetic modifiers (referred to as suppressors) that genetically interact with *fasn-1*, we performed an ethyl-methanesulfonate (EMS)-based forward genetic screen using the *fasn-1(g43ts)* mutant at a restrictive temperature of 25 °C. Leveraging the conditional lethality of *fasn-1(g43ts)*, we conducted a pilot screen involving over 88,000 haploid genomes (∼44,432 mutagenized F1s) at 25 °C (Fig.3A). To isolate the homozygous suppressor lines, embryonic lethality in each line was assessed for at least two generations, with testing conducted on at least 20 animals per generation. We confirmed 22 independent homozygous lines, all of which display consistently restored viability rates in each tested animal (Supplemental Fig.2B). Subsequently, the *fasn-1(g43ts)* allele was confirmed in each suppressor line by Sanger sequencing. The embryonic viability rate in each suppressor line was significantly restored when compared with the missense mutant alone, while the total number of F1 progenies was reduced, which suggests that the suppressor alleles may contribute to *C. elegans* fertility, or there are other essential alleles mutagenized in the suppressor background (Fig.3B-C, Supplemental Fig.2A-B). Our previous studies have shown that embryonic lethality in the *fasn-1* mutant is associated with the failure of the extracellular layer eggshell formation, which functions as extracellular protection to prevent embryos from osmotic and mechanical damage^19^. We therefore use the DAPI staining to assess the eggshell formation in each suppressor line. Nearly 100% embryos in the *fasn-1(g43ts)* exhibited DAPI penetration inside the cytosol (Fig.1F, 3E), while 50%-75% of embryos successfully blocked the DAPI in the tested suppressor lines (Fig.3D-E, Supplemental Fig.2C) at 25 °C. Overall, we established a forward genetic screen pipeline and successfully isolated suppressor lines, in which the putative suppressor variants alleviated the embryonic lethality and defective eggshell formation caused by the *fasn-1(g43ts)* mutant.

**Figure 3.**
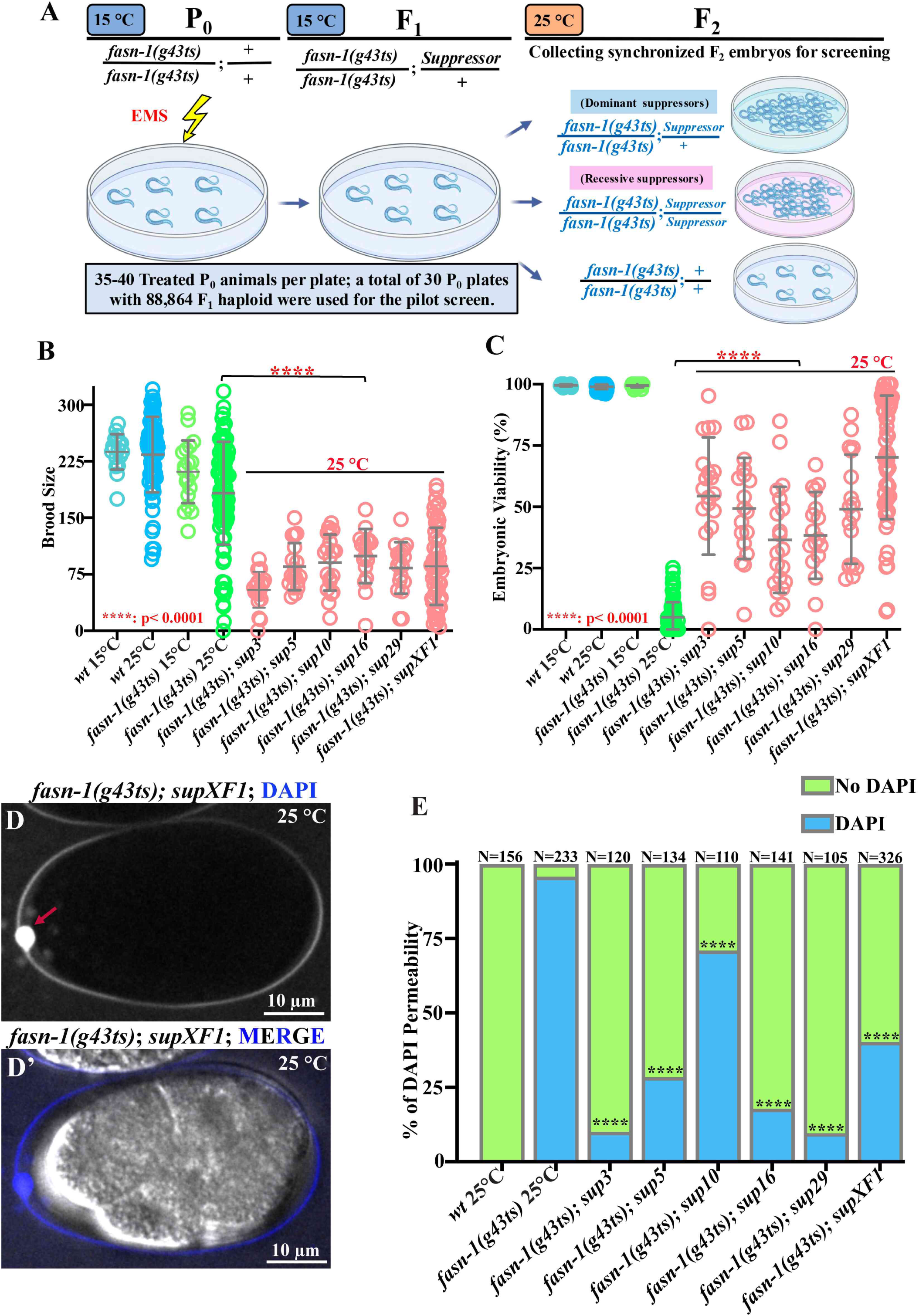
EMS-mediated forward genetic screens to identify genetic suppressors of the fasn-1(g43ts) mutant. (A) Workflow to conduct forward genetic screens to identify genetic modifiers that restore embryonic viability in *fasn-1(g43ts)* mutant. (B) Total brood size of the representative suppressor lines, wildtype, and *fasn-1(g43ts)* mutant only, over 60 hours post mid-L4 at different temperatures. (C) The percentage of embryonic viability was significantly restored in each suppressor line when compared with the *fasn-1(g43ts)* mutant only at 25 °C. (D-D’) Representative images of the DAPI staining assay showed only the polar body stained in the *fasn-1(g43ts)* suppressor line (red arrow in D). The merged images of DAPI and DIC were shown in panel D’. (E) Quantification of the embryos with DAPI staining in wildtype, *fasn-1(g43ts)* only, and the representative suppressor lines. Statistical significance was determined using a Chi-square test. ****: p< 0.0001

### Identification of the suppressor variants using genomic mapping and bioinformatic approaches

To identify the genetic modifiers responsible for restoring the viability in the suppressor lines, we combined two complementary approaches, including: (1) molecular inversion probes mapping (MIPs) for genetically mapping mutations in the candidate strains (a technique known as MIP-MAPping) ^22^, and (2) a customized bioinformatics pipeline to pinpoint the putative suppressor mutations from the whole-genome sequencing data (Fig.4A). MIP-MAP is a targeted genetic mapping technique that uses a strain, VC20019, from the Million Mutation Project^23^ containing MIP-verified single-nucleotide variants (SNVs) distributed across the *C. elegans* genome at approximately 1 Mb intervals^22^. To conduct this mapping, we introduced the *fasn-1(g43ts)* allele in VC20019 by CRISPR/Cas9 gene editing, and renamed this strain *fasn-1(g43ts)^MM^*. Then the suppressor line was crossed with the *fasn-1(g43ts)^MM^* reference strain, and the progeny were propagated for over ten generations to select for homozygosity of the suppressor mutation. Subsequently, the SNVs are sequenced, and the regions where the known SNV frequencies drop to zero are interpreted as loci harboring the suppressor mutation (Supplemental Fig.3A-B). Although this method is effective, it is both labor-intensive and time-consuming.

**Figure 4.**
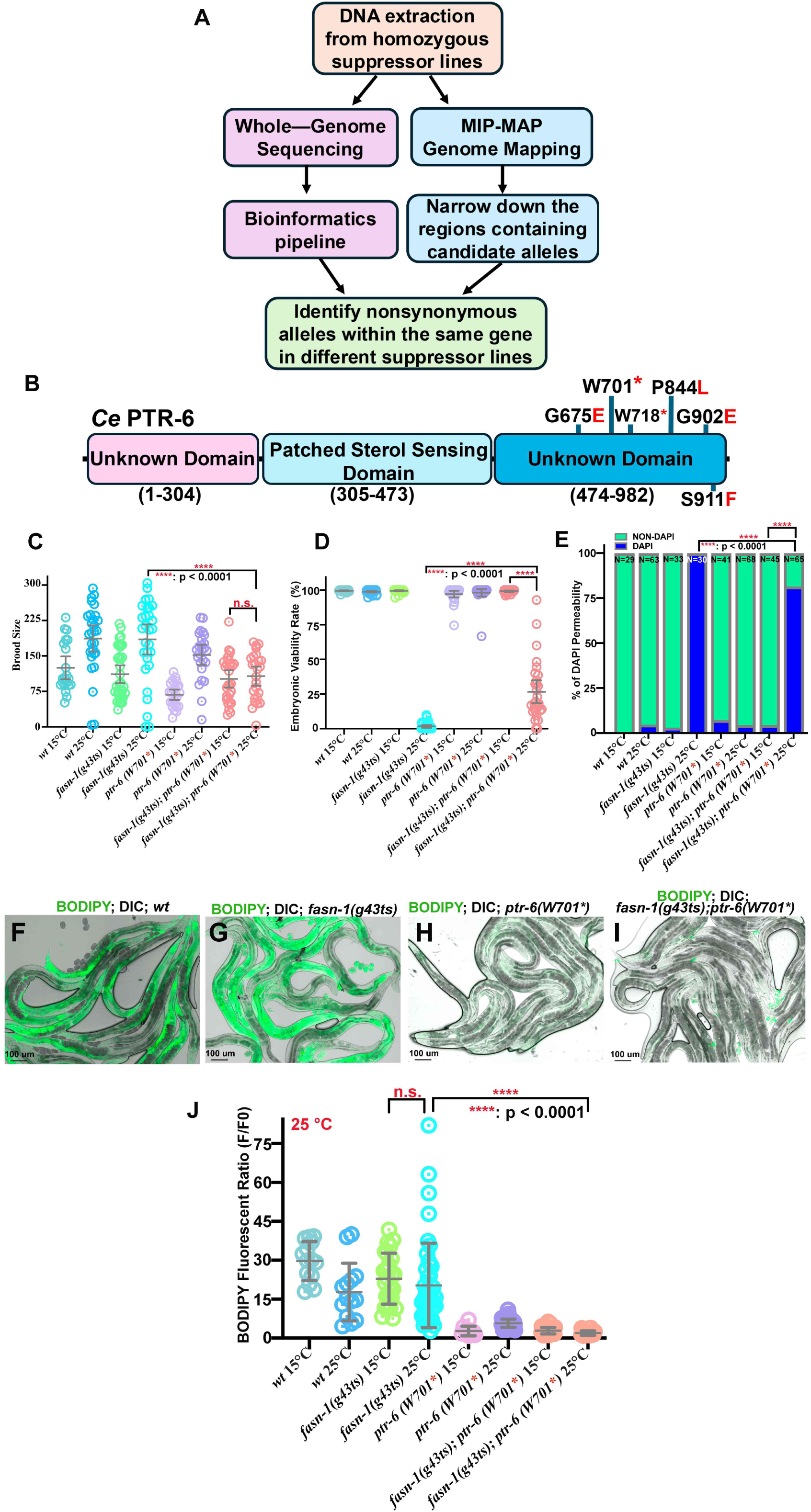
Identification of the genetic modifiers by whole-genome sequencing and a customized bioinformatic pipeline. (A) Workflow to identify the genetic modifiers from suppressor lines. (B) Schematic diagram of six suppressor alleles of *ptr-6*. (C-D) Total brood size and embryonic viability of wildtype, *fasn-1(g43ts)* only, *ptr-6(W791*)* suppressor alleles, and double mutants of *fasn-1(g43ts)* and *ptr-6(W791*)* allele at both 15 °C and 25 °C. (E) Quantification of the DAPI staining of embryos at different temperatures. Scale bars are indicated in each panel. (F-I) Representative images of the BODIPY-stained animals of wild-type, *fasn-1(g43ts)* only, *ptr-6(W791*)* suppressor alleles, and double mutants of *fasn-1(g43ts)* and *ptr-6(W791*)* at 25 °C. (J) Quantification of the BODIPY fluorescent intensity in the germline at different temperatures. Statistical significance was determined using One-Way ANOVA. ****: p< 0.0001. Scale bars are indicated in each panel of F-I.

In previous suppressor screens, we have identified both independent suppressor in the same gene as well as mutations in different genes within the same genetic or signaling pathway across independent suppressor lines^20, 24, 25^. Based on this observation, we developed a bioinformatics pipeline to prioritize candidate alleles that affect the same genes across independent suppressor lines. We filtered the variants with frameshift, stop gain, loss, missense, and in-frame indel variants. Genes with multiple independent variants across samples are prioritized for the downstream studies. This bioinformatic approach streamlines the identification of causative mutations, significantly reducing the time and labor required for MIP-MAP mapping (Fig.4A). To test the feasibility of this new strategy, we applied whole-genome data from all 22 suppressor lines to our bioinformatics pipeline. By integrating this with extensive gene function analysis, we identified six missense mutations in the *ptr-6* gene as one of our top suppressor candidates, which is involved in the Hedgehog signaling pathway (Fig. 4B). The *ptr-6* gene was also mapped in our MIP-MAP approaches on Chromosome II in one of the suppressor lines *fasn-1(g43ts); supXF1* (Supplemental Fig.3B), suggesting the feasibility and successful strategy to combine genomic mapping and bioinformatic pipeline to isolate the genetic variants.

### Hedgehog-related gene *ptr-6* genetically compensates for the embryonic lethality caused by *fasn-1(g43ts)*

All six identified suppressor alleles are in the C-terminal region of the PTR-6 protein, with two introducing premature stop codons (Fig.4B). The predicted PTR-6 structure from AlphaFold suggests that the residues were located at the C-terminus of different helical segments (Supplemental Fig.4A-F). To validate the role of *ptr-6* in suppression, we generated the *ptr-6(W701*)* allele in both wild-type and *fasn-1(g43ts)* backgrounds (Fig. 6B) using CRISPR/Cas9 gene editing. The homozygous *ptr-6(W701*)* animals exhibited nearly 100% embryonic viability, proper eggshell formation, and behaved as superficial wild-type at 25°C, suggesting that this missense mutation may not significantly disrupt the essential function of PTR-6 in *C. elegans* (Fig. 4C-E). However, the *ptr-6(W701*); fasn-1(g43ts)* double mutant showed a significant restoration of viability and defective eggshell formation, but reduced brood size compared to *fasn-1(g43ts)* alone at 25 °C (Fig.4C-E).

To further assess whether loss-of-function of *ptr-6* contributes to the suppression of *fasn-1*, we fed *ptr-6*(RNAi) bacteria to both wild-type and *fasn-1(g43ts)* mutants to globally knock down the *ptr-6* expression at 25 °C. We first fed *ptr-6(RNAi)* to endogenously tagged PTR-6::AID::GFP animals as a fluorescent reporter to assess the knockdown efficiency and specificity of *ptr-6* RNAi. The fluorescent intensity of PTR-6::AID::GFP in the somatic tissue, including the head excretory cell, where the PTR-6 is highly expressed, was dramatically reduced compared with the RNAi negative control, suggesting that *ptr-6* RNAi effectively knocked down the *ptr-6* gene expression *in vivo* (Supplemental Fig.5A-D). We then depleted *ptr-6* by RNAi in *fasn-1(g43ts)* mutants at 25 °C (Supplemental Fig.5E). The *ptr-6(RNAi)* significantly increased embryonic viability in *fasn-1(g43ts)* tested animals compared to RNAi negative control (Supplemental Fig.5E), suggesting that *ptr-6* reduction-of-function indeed mitigates *fasn-1*-associated embryonic defects.

**Figure 5.**
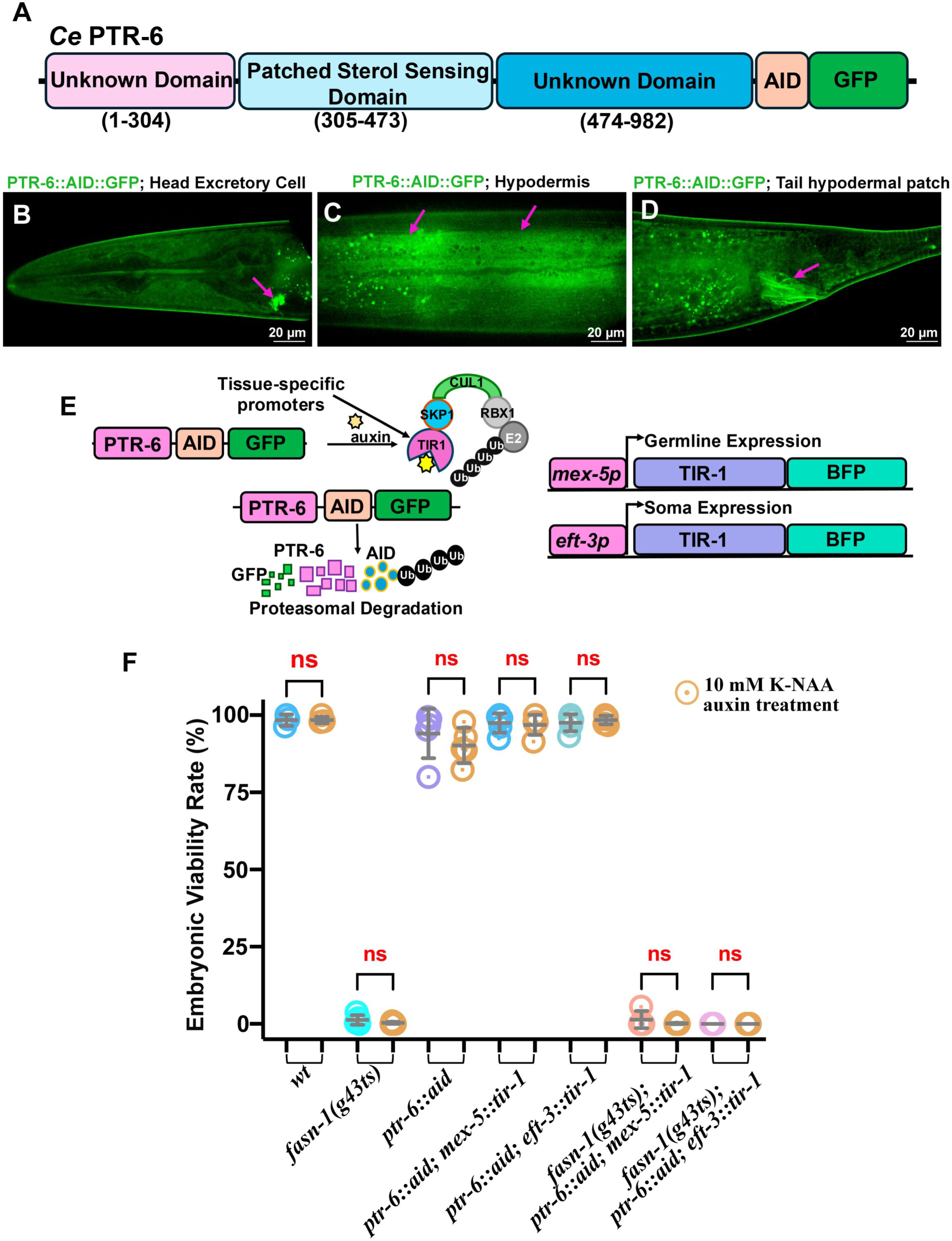
PTR-6::AID::GFP is expressed in multiple tissues. (A) A degron and GFP cassette was inserted at the 3′ end of the *ptr-6* coding sequence using CRISPR/Cas9 genome editing. (B-D) PTR-6::AID::GFP is expressed in multiple tissues, including the head excretory cell (magenta arrow in B), hypodermis (magenta arrows in C), and tail hypodermal patch area (magenta arrow in D). (E) Two transgenic TIR-1::BFP::AID lines driven by different promoters, including soma-specific *eft-3* promoter and the germline-specific promoter *mex-5,* were crossed with the PTR-6::AID::GFP line to activate the tissue-specific degradation of PTR-6. (F) Embryonic viabilities of *fasn-1(g43ts)* were not affected when treated with 10 mM auxin K-NAA in each PTR-6::AID::GFP strain. The orange circles represent those animals treated with auxin.

The *ptr-6* is the *C. elegans* ortholog of human *PTCHD3*, which is predominantly expressed in epidermal tissues and is predicted to regulate membrane integrity, permeability, and molting processes in *C. elegans* ^26, 27^. We therefore hypothesized that the observed suppression is linked to changes in membrane integrity or cuticle permeability. To test this hypothesis, we performed *in vivo* staining with BODIPY, a lipophilic dye that penetrates the cuticles to label lipid droplets and neutral lipids in various tissues or cells, thereby allowing for the assessment of cuticle integrity and the visualization of overall lipid content *in vivo*. The *fasn-1(g43ts)* mutant showed moderately increased BODIPY staining relative to wild-type, though the difference was not statistically significant, suggesting either altered lipid storage in the mutant or feenhanced dye penetration in the *fasn-1(g43ts)* background (Fig.4F-G, J). In contrast, *ptr-6(W701*)* and *ptr-6(W701*); fasn-1(g43ts)* double mutants showed a significant reduction of the BODIPY penetration relative to both *fasn-1(g43ts)* and wild-type controls (Fig.4H-J).

These results suggest two possible hypotheses: (1) BODIPY penetration is reduced in the *ptr-6(W701*)* mutant due to altered cuticle or membrane integrity, or (2) loss of *ptr-6* affects lipid synthesis or metabolism, potentially compensating for the lipo-toxicity associated with *fasn-1(g43ts)*. The presence of DMSO in the BODIPY staining buffer, which can dehydrate the animals and alter membrane fluidity, typically results in the animals showing temporal immobility or slow locomotion (Video 1). Intriguingly, *ptr-6* mutants retained faster locomotion than wild-type and *fasn-1* mutants during staining at both 15°C and 25 °C. Quantitative analysis confirmed that *ptr-6(W701*)* and *ptr-6(W701*)*; *fasn-1(g43ts)* double mutants were significantly dynamic than the wild-type and *fasn-1(g43ts)* mutants only (Video 1; Supplemental Fig.6). This observation is significantly enhanced when the animals are maintained at a higher temperature of 25 °C. These observations suggest that the loss of *ptr-6* may increase membrane or cuticle permeability, preventing BODIPY penetration, which is consistent with previous findings that the *ptr-6* null mutant displays increased resistance to ethanol due to altered membrane integrity and permeability^26^. Given the importance of fatty acid synthase in lipid metabolism and physiology, our observation does not rule out the possibility that this genetic suppression was not associated with lipid metabolic imbalance. However, given the complexity of lipid metabolism, we decided to focus on genetic screens and permeability assays in this study and will investigate the lipid metabolic aspect in a separate study.

### Tissue-specific degradation of PTR-6 does not suppress the embryonic lethality of *fasn-1(g43ts)*

To better understand the role of PTR-6 in suppressing the embryonic lethality of the *fasn-1(g43ts)* mutant, we used an auxin-inducible degradation system (AID) to degrade PTR-6 in somatic tissues or the germ line ^28^. We inserted a cassette with degron and GFP coding sequences at the *ptr-6* C-terminus using CRISPR/Cas9 genome editing (named PTR-6::AID::GFP), which enables us to visualize and quantify the degradation efficiency (Fig.5A-E). We used the *Peft-3* and *Pmex-5* to drive the somatic and germline-specific AID strains, respectively, and activated degradation when the animals were exposed to 10 mM 1-naphthalenacetic acid (1-NAA) ^29^. The GFP fluorescence intensities in the induced animals were significantly reduced compared to those in the control (Supplemental Fig.6A-G). The *Peft-3::tir-1::BFP::AID* led to an approximate 1.6-fold reduction of fluorescent intensity of PTR-6::AID::GFP in the head and other somatic tissues (Supplemental Fig.6A-G), suggesting the degradation of PTR-6::AID::GFP was affected in the somatic tissues. We did not observe any fluorescent intensity of PTR-6::AID::GFP in the germline tissue or early embryos, suggesting that PTR-6 may have no or low expression in the reproductive tissues. Therefore, we were unable to assess the degradation of PTR-6 in the *Pmex-5::tir-1::BFP::AID* background.

To assess the tissue-specific suppression of PTR-6 in *fasn-1(g43ts)*, we crossed the *fasn-1(g43ts)* allele into each degron strain. We exposed L4 animals to either regular food as a control or 10 mM K-NAA, then embryonic viability rate was determined 0-60 hours post-L4 at 25 °C (Fig.5F). Unexpectedly, the embryonic viability of the *fasn-1(g43ts)* mutant was not restored in either somatic or germline tissue-specific PTR-6::AID::GFP strains (Fig.5F). These findings suggested that the AID system might not sufficiently degrade PTR-6 below the basal level to compromise the embryonic lethality in the *fasn-1(g43ts)* mutant.

## Discussion

Our study identified and validated the hedgehog signaling-associated molecule *ptr-6* as a genetic suppressor of the fatty acid synthase gene *fasn-1* in *C. elegans*, revealing a previously unknown connection between fatty acid synthesis and Hedgehog-like signaling pathways during embryonic development. As the central rate-limiting enzyme in *de novo* lipogenesis, FASN-1 is essential to synthesize long-chain fatty acids that are critical to membrane biogenesis and cellular signaling production. We found that dysfunction of FASN-1 in *C. elegans* disrupts the formation of the extracellular permeability barrier, leading to defects in embryogenesis and early development. By conducting EMS-mediated forward genetic screens, we identified six *ptr-6* mutations that significantly restore embryonic viability and rescue permeability defects, suggesting that *ptr-6* acts as a genetic compensator for mitigating the consequences of FASN-1 loss-of-function and potential lipid biosynthetic deficiency.

PTR-6 is a patched-domain protein, which is homologous to human PTCHD3. PTR-6 is predominantly expressed in epidermal tissues, where it is predicted to regulate membrane integrity and cuticle permeability^26^. The genetic suppression of loss-of-function *ptr-6* in the *fasn-1(g43ts)* mutant in the context of permeability suggested that the PTR-6 protein contributes, directly or indirectly, to maintaining cuticular or cellular membrane properties that are particularly sensitive to proper fatty acid synthesis or lipid homeostasis. One plausible hypothesis is that PTR-6 could restrain the flexibility or permeability of the embryonic eggshell and somatic membranes, while losing PTR-6 function might reduce these constraints and enhance membrane fluidity to compensate for defects in lipid composition in the FASN-1 mutant. This hypothesis is supported by our BODIPY staining data, where *ptr-6* and *fasn-1* double mutant significantly reduced the penetration of the staining molecule, which reflects a change in membrane barrier integrity or fluidity. Moreover, the faster locomotion observed in *ptr-6* mutants during staining suggests broader changes in membrane dynamics that could improve stress resilience under high-temperature or chemical perturbation.

Previous studies have shown that Hedgehog-related pathways mediated by patched-domain proteins respond to sterol or lipid cues and modulate gene expression associated with proliferation, differentiation, extracellular matrix formation, and stress responses^30–33^ . The *C. elegans* PTR family may play analogous roles by potentially linking lipid availability to the transcriptional control of stress-responsive genes and extracellular matrix synthesis ^27, 31–33^. Our findings are also consistent with a recent study showing that the C-terminal fragment of FASN (FASN-CTF) influences the stress-responsive signaling pathways, including the components of Hedgehog-like signaling pathways, particularly under conditions of lipid imbalance ^34^. Additionally, our data suggest that loss of *ptr-6* may also directly affect lipid metabolism or synthesis. Previous studies suggest that the *C. elegans* PTR proteins may influence lipid trafficking, storage, or utilization in epidermal tissues, which in turn affects germline development, eggshell biogenesis, or extracellular matrix formation^31–33^. Given the connection between somatic tissues and the germline in *C. elegans*, changes in epidermal or hypodermal lipid composition could modulate the supply of fatty acids to developing embryos, partially compensating for reduced FASN activity^12^. Further lipidomic and transcriptomic analyses will be needed to clarify whether and how *ptr-6* compromises the imbalanced or disrupted fatty acid composition in the *fasn-1* mutant.

In conclusion, our work identified a previously uncharacterized genetic compensation mechanism that alleviates the embryonic defects caused by *fasn-1* mutants, potentially involving Hedgehog-like signaling in *C. elegans*. Dysfunction of *fasn-1* caused eggshell defects and embryonic lethality, whereas loss of *ptr-6* restores permeability and viability, possibly by mitigating defects in membrane flexibility, extracellular matrix formation, or lipid imbalance. Finally, our study established a genetic screen pipeline for probing the intersection of lipid metabolism and developmental signaling *in vivo*. Future studies will integrate genetic, biochemical, and lipidomic approaches to further investigate the mechanisms of how PTR-6 and other Patch-domain proteins compensate for metabolic deficiencies and maintain developmental homeostasis in the *fasn-1* mutant.

## Materials and Methods

### *C. elegans* strains used in this study

All *C. elegans* strains were maintained on MYOB plates with the OP50 bacteria as a food source, unless otherwise specified in other sections. Strain information is listed in the supplemental information.

### EMS-mediated forward genetic screen

The early L4 hermaphrodites of XFB2 *fasn-1(g43ts)* were synchronized and washed three times in M9 and soaked in 48 mM Ethyl methane sulfonate (EMS) solution for four hours. The EMS-treated animals were rinsed three times in M9 buffer and were transferred to multiple fresh 100 mm MYOB plates with OP50 on one side. The animals were allowed to recover for up to 4 hours before being picked and placed individually on 100 mm MYOB plates with fresh OP50. Only the recovered late L4 or young adult animals that were able to crawl across the plates to the OP50 food were transferred to the fresh plates. A total of 30 MYOB plates with 25-40 mid-L4 (a.k.a. P0s) on each were incubated at 15 °C. Gravid F1 adults were bleached, and F2 embryos were collected after hypochlorite treatment. The F2 embryos were shaken in a 15 ml Falcon tube with M9 buffer overnight, and hatched larvae were grown on OP50 plates at 15 °C until the L4 stage. Subsequently, all plates were incubated at 25 °C for a week to screen suppressor candidates. Their progeny was screened for viable larvae. ∼88,000 mutagenized haploid genomes were scored in the screens.

### RNAi treatment

The RNAi feeding constructs were selected from either the Ahringer or Vidal libraries. The activated RNAi bacteria culture was shaken for five hours until the log phase was reached. The bacterial stock was resuspended in fresh LB broth containing 1 mM IPTG and 25 μg/ml carbenicillin before being spread on MYOB plates. The medium plates were then incubated for 12-14 hours at 15 °C to enhance the activation of the RNAi plasmid expression before being fed to the animals. The seeded RNAi plates were also freshly prepared the day prior to treatment. To deplete the target gene *ptr-6*, mid-L4 hermaphrodites were picked onto plates with the IPTG-induced bacteria. Animals were grown on RNAi plates at 25°C for 36-60 hours for the embryonic lethality and other assays.

### Embryonic viability assay

Single mid-L4 hermaphrodites were picked onto 35 mm MYOB plates seeded with 10 μl of fresh OP50 or RNAi bacteria and allowed to lay eggs for 24 hours post mid-L4 at 25°C. The hermaphrodites were transferred to a newly seeded 35 mm MYOB plate to lay eggs for an additional 24 hours. The transfer was repeated three times before the adult hermaphrodites were flamed from the plate. Twenty-four hours after removing the hermaphrodites, the viable larvae were counted for embryonic viability. The total brood sizes were determined at 72 hours. Embryonic viability rate = (the number of hatched larvae / the total brood size) *100%.

### CRISPR design

All CRISPR/Cas9 editing was generated into the Bristol N2 strain as the wild type, unless otherwise indicated. The crRNAs were synthesized from Horizon Discovery, along with tracrRNA. The repair template and oligos were purchased from Integrated DNA Technologies (IDT). Approximately 20-30 young gravid animals were injected with the CRISPR/Cas9 injection mix. Detailed sequence information of CRISPR design was listed below in 3: (Capital letters represent the ORF or exon sequence, lowercase letters indicate the sequence from the intron). Detailed information is listed in the supplementary information.

### Auxin-inducible treatment in the degron strains

Auxin 1-naphthaleneacetic potassium salt (K-NAA) was purchased from PhytoTech (#N610). A 1 M stock solution of K-NAA was prepared in water and added to MYOB medium to achieve a final concentration of 4-10 mM auxin. To efficiently degrade the PTR-6 protein, mid-L4 hermaphrodites were picked onto auxin plates. Animals were grown on the plates at 20°C for 24 hours for the degradation efficiency test, and 60 hours for the brood size assay.

### Microscopy

All imaging was performed on a spinning disk confocal system that uses a Nikon 60X 1.2 NA water or oil objectives, a Hamamatsu C15440 ORCA-Fusion BT Digital camera, and a Yokogawa CSU-X1 confocal scanner unit. Nikon’s NIS imaging software was applied to capture the images. The image data were processed using the ImageJ/FIJI Bio-formats plugin (National Institutes of Health)^35, 36^.

### MIP-MAP preparation and bioinformatic analysis

*fasn-1(g43ts^MM^)* males were mated with the homozygous hermaphrodites of each suppressor line (Fig. S2A). We carefully pooled suppressed F2 progeny from the cross and allowed the F2 population to expand for ten generations for MIP-MAP analysis. At least one generation of self-recombination is sufficient to distribute the MIP-MAP single-nucleotide polymorphism (SNPs) (also referred to as molecular probes) into the suppressor line background and provide a high molecular resolution to identify the mutated regions. Candidate mutations (defined as novel, homozygous, and nonsynonymous) were identified by whole-genome sequencing as described previously^37, 38^. Briefly, sequencing libraries were constructed by Invitrogen Pure Link Genomic DNA Mini Kit (Ref # K1820-01) with genomic DNA from homozygous suppressor-bearing strains. The libraries were pooled and sequenced on a HiSeq 3000 instrument (Illumina, San Diego, CA) to at least 20-fold coverage. Variants were identified using a pipeline as described in our previous studies^38, 39^. Mapping loci for suppressors was identified using molecular inversion probes (MIPs) to single-nucleotide polymorphisms (SNPs), as described previously^38–40^. Briefly, suppressor-bearing strains were mated to SNP mapping strain VC20019, which had been engineered via CRISPR to contain the *fasn-1(g43ts)* mutation. F1 cross-progeny were allowed to self-fertilize, and a minimum of 50 homozygous F2 progeny were pooled for the construction of MIP libraries. SNP allele frequencies were determined using a custom script and plotted with R to delimit the mapping interval as previously described^38, 39^.

For the customized bioinformatic pipeline, we used BWA-MEM2 to map and align paired-end reads to the *Caenorhabditis elegans* reference genome (accession PRJNA13758.WS292), Picard v.1.93 MarkDuplicates to identify and flag PCR duplicates, DeepVariant v.0.8.0 to generate individual-level gVCF files from the aligned sequence data, and GLnexus v1.1.3 to jointly call the variants. Phred-scaled quality score, allele depths were used for variant quality control. Variant annotation was added from PRJNA13758.WS245.annotations using bcftools plugin version 1.19. Variants predicted to be frameshift, stop gain, stop loss, essential splice site, missense, and in-frame indel variants were selected. Genes with multiple variants identified in different samples were prioritized. The detailed sequencing and bioinformatic analysis of the independent suppressor lines will be available upon request.

### Data and statistical analyses

Statistical significance for other assays was determined by p-value from an unpaired 2-tailed t-test, one-way ANOVA, or Chi-squared test. P-values: n.s. = not significant; * = <0.05; ** = <0.01, *** = <0.001; **** = <0.0001. Both the Shapiro-Wilk and Kolmogorov-Smirnov Normality test indicated that all data follow normal distributions.

## Acknowledgments

We thank the Caenorhabditis Genetics Center, which is funded by the National Institutes of Health Office of Research Infrastructure Programs (P40OD010440), for providing strains for this study. We also thank Dr. Kota Mizumoto for generously sharing the plasmid pSM.GFPnovo2^41^. We thank all members of the UF Worm clubs for providing feedback and suggestions to our investigations. In Memoriam of Dr. Andy Golden, who provided tremendous scientific support to initiate the ideas for this study.

## Funding

The project was supported by an NIH Pathway to Independence Award (K99/R00), 1K99 GM145224-01(X.F.B.). National Institute of General Medical Sciences/National Institutes of Health (R00GM145224 to X.F.B.), and the UF Startup fund for the Bai lab. The project was, in part, supported by the National Institute of Diabetes and Digestive and Kidney Diseases/National Institutes of Health Intramural Research funding (to H.E.S.) and the National Human Genome Research Institute/ /National Institutes of Health (R00HG011490 to X.F.).

---

**Supplementary Figure 1. FASN-1 is expressed in multiple *C. elegans* tissues.** (A) Schematic diagram of both N- and C-terminus endogenously tagged FASN-1 reporters, including GFP::FASN-1 and FASN-1::RFP. (B-C’’) Overlap of both GFP::FASN-1 and FASN-1::RFP expression in excretory tissues (B-B’’) and gonadal sheath cells (C-C’’). Scale bars are indicated in each panel.

**Supplementary Figure 2. Embryonic viability and eggshell formation were restored in suppressor lines.** (A-B) Total brood size (A) and the percentage of embryonic viability (B) of each isolated suppressor line, wild type, and the *fasn-1(g43ts)* mutant only over 60 hours post mid-L4 at non-permissive temperature at 25 °C. (C) Quantification of the embryos with DAPI staining in wild type, *fasn-1(g43ts)* only, and indicating suppressor lines. Statistical significance was determined using a One-Way ANOVA in panel A-B and a Chi-square test in panel C. ****: p< 0.0001, * p<0.05; ** p< 0.01; *** p<0,001; **** p< 0.0001.

**Supplementary Figure 3. Identification of the genetic modifiers by MIP-MAP genome mapping and whole-genome sequencing.** (A) Diagram of the MIP-MAP workflow to map the SNVs in the suppressor line. (B-F) The read frequency of the VC20019-specific SNVs across the genome of the pooled F2 progeny from *fasn-1(g43ts)*; *supXF1* and *fasn-1(g43ts)^MM^* cross. (B) An identical candidate mutation-associated interval was identified on Chromosome II in *fasn-1(g43ts)*; *supXF1*. The strategy graphic was generated with BioRender.com.

**Supplementary Figure 4. The missense residues of PTR-6 were located on the unknown C-terminus domain.** (A-F) The structure of PTR-6 from Alphafold indicated that all six missense mutations were on the C-terminal of different helices (highlighted by red arrows. Distinguished colors represent the confidence rate of the protein structure, including dark blue-Very high (pLDDT > 90); cyan-Confident (90 > pLDDT > 70); yellow-Low (70 > pLDDT > 50); orange-Very low (pLDDT < 50).

**Supplementary Figure 5. ptr-6 RNAi is efficient and specific to reduce the expression of ptr-6::aid::gfp in the somatic tissue.** (A-B) PTR-6::AID::GFP was strongly expressed in the head excretory cell (magenta arrow in A-B). (C) Knock-down *ptr-6* gene expression by RNAi significantly reduced the fluorescent signals of the PTR-6::AID::GFP cassette in the excretory cell (magenta arrow in C). (D) Quantification of fluorescent intensity of head PTR-6::AID::GFP with OP50, *ctrl*(RNAi), and *ptr-6*(RNAi) treatment. (E) *ptr-6*(RNAi) treatment significantly restored the embryonic viability in the *fasn-1(g43ts)* mutant at the restrictive temperature. The RNAi experiments were completed with three independent replicates. Scale bars are indicated in each panel. Statistical significance was determined using a One-Way ANOVA. P values: ****: p< 0.0001.

**Supplementary Figure 6. Locomotion of the animals treated with BODIPY staining.** (A) Quantification of body bend frequency of wild type, *fasn-1(g43ts)* only, *ptr-6(W701*)* only, and *fasn-1(g43ts)*; *ptr-6(W701*)* double mutants after BODIPY staining. The body bends were measured over a 60-second recording period at both 15 °C and 25 °C.

**Supplementary Figure 7. Tissue-specific degradation of PTR-6 significantly reduced the fluorescence of the PTR-6::AID::GFP cassette in each tissue expressing TIR-1::mTagBFP2::AID.** (A–F′′) PTR-6::AID::GFP localizes to somatic tissues, such as head excretory cell, hypodermis, and tail hypodermal patch (magenta arrows in A, A′′, C, C’’, and E, E’’). (A-F’’) TIR-1::mTagBFP2::AID driven by the somatic-specific promoters *eft-3* is strongly expressed in the nuclei of all or most somatic tissues (A’-A’’, C’-C’’, and E’-E’’). Fluorescent signals of both PTR-6::AID::GFP and TIR-1::mTagBFP2::AID driven by *eft-3* promoter at head excretory cell, hypodermis, and tail hypodermal patch are significantly reduced when animals were treated with 10 mM auxin K-NAA (magenta arrows in B-B′′, D-D’’, and F-F’’). (G) Quantification of the fluorescent signals of PTR-6::AID::GFP under control and 10 mM K-NAA treatment. P-values: *: p< 0.05 (t-test).

**Video 1. Behavioral analysis of animals treated with BODIPY staining.** Representative locomotion videos of wild type, *fasn-1(g43ts)* only, *ptr-6(W701*)* only, and *fasn-1(g43ts)*; *ptr-6(W701*)* double mutants after BODIPY staining. The videos were recorded over a 60-second period at both 15 °C and 25 °C.

## Bibliography

1. Ameer F, Scandiuzzi L, Hasnain S, Kalbacher H, Zaidi N. De novo lipogenesis in health and disease. Metabolism. 2014;63(7):895–902. Epub 20140412. doi: 10.1016/j.metabol.2014.04.003. PubMed PMID: 24814684.

2. Gonzalez-Bohorquez D, López IMG, Jaeger BN, Pfammatter S, Bowers M, Semenkovich CF, Jessberger S. FASN-dependent de novo lipogenesis is required for brain development. P Natl Acad Sci USA. 2022;119(2). doi: ARTN e2112040119 10.1073/pnas.2112040119. PubMed PMID: WOS:000768580200004.

3. Sturmey RG, Reis A, Leese HJ, McEvoy TG. Role of fatty acids in energy provision during oocyte maturation and early embryo development. Reprod Domest Anim. 2009;44 Suppl 3:50–8. doi: 10.1111/j.1439-0531.2009.01402.x. PubMed PMID: 19660080.

4. Wathes DC, Abayasekara DR, Aitken RJ. Polyunsaturated fatty acids in male and female reproduction. Biol Reprod. 2007;77(2):190–201. Epub 20070418. doi: 10.1095/biolreprod.107.060558. PubMed PMID: 17442851.

5. Lin HP, Cheng ZL, He RY, Song L, Tian MX, Zhou S, Groh BS, Liu WR, Ji MB, Ding C, Shi YH, Guan KL, Ye D, Xiong Y. Destabilization of Fatty Acid Synthase by Acetylation Inhibits Lipogenesis and Tumor Cell Growth. Cancer Res. 2016;76(23):6924–36. doi: 10.1158/0008-5472.Can-16-1597. PubMed PMID: WOS:000389435800018.

6. Maier T, Leibundgut M, Ban N. The crystal structure of a mammalian fatty acid synthase. Science. 2008;321(5894):1315–22. doi: 10.1126/science.1161269. PubMed PMID: 18772430.

7. Wang YX, Voy BJ, Urs S, Kim S, Soltani-Bejnood M, Quigley N, Heo YR, Standridge M, Andersen B, Dhar M, Joshi R, Wortman P, Taylor JW, Chun J, Leuze M, Claycombe K, Saxton AM, Moustaid-Moussa N. The human fatty acid synthase gene and de novo lipogenesis are coordinately regulated in human adipose tissue. J Nutr. 2004;134(5):1032–8. PubMed PMID: WOS:000221423000007.

8. Chirala SS, Chang H, Matzuk M, Abu-Elheiga L, Mao JQ, Mahon K, Finegold M, Wakil SJ. Fatty acid synthesis is essential in embryonic development:: Fatty acid synthase null mutants and most of the heterozygotes die. P Natl Acad Sci USA. 2003;100(11):6358–63. doi: 10.1073/pnas.0931394100. PubMed PMID: WOS:000183190700015.

9. Fhu CW, Ali A. Fatty Acid Synthase: An Emerging Target in Cancer. Molecules. 2020;25(17). doi: ARTN 3935 10.3390/molecules25173935. PubMed PMID: WOS:000569618300001.

10. Vanauberg D, Schulz C, Lefebvre T. Involvement of the pro-oncogenic enzyme fatty acid synthase in the hallmarks of cancer: a promising target in anti-cancer therapies. Oncogenesis. 2023;12(1). doi: ARTN 16 10.1038/s41389-023-00460-8. PubMed PMID: WOS:000952323000001.

11. Landrock D, Atshaves BP, McIntosh AL, Landrock KK, Schroeder F, Kier AB. Acyl-CoA Binding Protein Gene Ablation Induces Pre-implantation Embryonic Lethality in Mice. Lipids. 2010;45(7):567–80. doi: 10.1007/s11745-010-3437-9. PubMed PMID: WOS:000279695400001.

12. Starich TA, Bai XF, Greenstein D. Gap junctions deliver malonyl-CoA from soma to germline to support embryogenesis in. Elife. 2020;9. doi: ARTN e58619 10.7554/eLife.58619. PubMed PMID: WOS:000563771500001.

13. Huelgas-Morales G, Sanders M, Mekonnen G, Tsukamoto T, Greenstein D. Decreased mechanotransduction prevents nuclear collapse in a laminopathy. P Natl Acad Sci USA. 2020;117(49):31301–8. doi: 10.1073/pnas.2015050117. PubMed PMID: WOS:000598990900013.

14. Lynn DA, Dalton HM, Sowa JN, Wang MC, Soukas AA, Curran SP. Omega-3 and-6 fatty acids allocate somatic and germline lipids to ensure fitness during nutrient and oxidative stress in. P Natl Acad Sci USA. 2015;112(50):15378–83. doi: 10.1073/pnas.1514012112. PubMed PMID: WOS:000366404200054.

15. Killian DJ, Hubbard EJA. germline patterning requires coordinated development of the somatic gonadal sheath and the germ line. Dev Biol. 2005;279(2):322–35. doi: 10.1016/j.ydbio.2004.12.021. PubMed PMID: WOS:000227487200004.

16. Robinson-Thiewes S, Dufour B, Martel PO, Lechasseur X, Brou AAD, Roy V, Chen Y, Kimble J, Narbonne P. Non-autonomous regulation of germline stem cell proliferation by somatic MPK-1/MAPK activity in C. elegans. Cell Rep. 2021;35(8):109162. doi: 10.1016/j.celrep.2021.109162. PubMed PMID: 34038716; PMCID: PMC8182673.

17. Turmel-Couture S, Martel PO, Beaulieu L, Lechasseur X, Dzuna LVF, Narbonne P. Bidirectional transfer of a small membrane-impermeable molecule between the intestine and germline. J Biol Chem. 2024;300(12). doi: ARTN 107963 10.1016/j.jbc.2024.107963. PubMed PMID: WOS:001372312700001.

18. Watts JL, Browse J. Dietary manipulation implicates lipid signaling in the regulation of germ cell maintenance in. Dev Biol. 2006;292(2):381–92. doi: 10.1016/j.ydbio.2006.01.013. PubMed PMID: WOS:000236947900009.

19. Bai X, Woodbury D, Golden A. The fasn-1(g14ts) allele is a Gly1830Arg missense mutation in C. elegans FASN-1. MicroPubl Biol. 2020;2020. Epub 20200505. doi: 10.17912/micropub.biology.000244. PubMed PMID: 32550489; PMCID: PMC7252273.

20. Jaramillo-Lambert A, Fuchsman AS, Fabritius AS, Smith HE, Golden A. Rapid and Efficient Identification of Legacy Mutations Using Hawaiian SNP-Based Mapping and Whole-Genome Sequencing. G3-Genes Genom Genet. 2015;5(5):1007–19. doi: 10.1534/g3.115.017038. PubMed PMID: WOS:000354262000027.

21. Rappleye CA, Tagawa A, Le Bot N, Ahringer J, Aroian RV. Involvement of fatty acid pathways and cortical interaction of the pronuclear complex in Caenorhabditis elegans embryonic polarity. BMC Dev Biol. 2003;3:8. Epub 20031003. doi: 10.1186/1471-213X-3-8. PubMed PMID: 14527340; PMCID: PMC270048.

22. Mok CA, Au V, Thompson OA, Edgley ML, Gevirtzman L, Yochem J, Lowry J, Memar N, Wallenfang MR, Rasoloson D, Bowerman B, Schnabel R, Seydoux G, Moerman DG, Waterston RH. MIP-MAP: High-Throughput Mapping of Temperature-Sensitive Mutants via Molecular Inversion Probes. Genetics. 2017;207(2):447–63. doi: 10.1534/genetics.117.300179. PubMed PMID: WOS:000412232600005.

23. Thompson O, Edgley M, Strasbourger P, Flibotte S, Ewing B, Adair R, Au V, Chaudhry I, Fernando L, Hutter H, Kieffer A, Lau J, Lee N, Miller A, Raymant G, Shen B, Shendure J, Taylor J, Turner EH, Hillier LW, Moerman DG, Waterston RH. The million mutation project: A new approach to genetics in. Genome Res. 2013;23(10):1749–62. doi: 10.1101/gr.157651.113. PubMed PMID: WOS:000325202100017.

24. Bai XF, Smith HE, Golden A. Identification of genetic suppressors for a BSCL2 lipodystrophy pathogenic variant in Caenorhabditis elegans. Dis Model Mech. 2024;17(6). doi: ARTN dmm050524 10.1242/dmm.050524. PubMed PMID: WOS:001267657000004.

25. Bai XF, Smith HE, Romero LO, Bell B, Vásquez V, Golden A. A mutation in F-actin polymerization factor suppresses the distal arthrogryposis type 5 PIEZO2 pathogenic variant in Caenorhabditis elegans. Development. 2024;151(4). doi: ARTN dev202214 10.1242/dev.202214. PubMed PMID: WOS:001166879700001.

26. Choi MK, Son S, Hong M, Choi MS, Kwon JY, Lee J. Maintenance of Membrane Integrity and Permeability Depends on a Patched-Related Protein in. Genetics. 2016;202(4):1411-+. doi: 10.1534/genetics.115.179705. PubMed PMID: WOS:000373959100015.

27. Zugasti O, Rajan J, Kuwabara PE. The function and expansion of the Patched- and Hedgehog-related homologs in. Genome Res. 2005;15(10):1402–10. doi: 10.1101/gr.3935405. PubMed PMID: WOS:000232436800011.

28. Zhang L, Ward JD, Cheng Z, Dernburg AF. The auxin-inducible degradation (AID) system enables versatile conditional protein depletion in C. elegans. Development. 2015;142(24):4374–84. Epub 20151109. doi: 10.1242/dev.129635. PubMed PMID: 26552885; PMCID: PMC4689222.

29. Ashley GE, Duong T, Levenson MT, Martinez MAQ, Johnson LC, Hibshman JD, Saeger HN, Palmisano NJ, Doonan R, Martinez-Mendez R, Davidson BR, Zhang W, Ragle JM, Medwig-Kinney TN, Sirota SS, Goldstein B, Matus DQ, Dickinson DJ, Reiner DJ, Ward JD. An expanded auxin-inducible degron toolkit for Caenorhabditis elegans. Genetics. 2021;217(3). doi: 10.1093/genetics/iyab006. PubMed PMID: 33677541; PMCID: PMC8045686.

30. Nguyen TD, Truong ME, Reiter JF. The Intimate Connection Between Lipids and Hedgehog Signaling. Front Cell Dev Biol. 2022;10. doi: ARTN 876815 10.3389/fcell.2022.876815. PubMed PMID: WOS:000815144100001.

31. Sundaram MV, Pujol N. The Caenorhabditis elegans cuticle and precuticle: a model for studying dynamic apical extracellular matrices in vivo. Genetics. 2024;227(4). doi: 10.1093/genetics/iyae072. PubMed PMID: WOS:001266565500001.

32. 32. Cohen JD, del Castillo CEC, Serra ND, Kaech A, Spang A, Sundaram M. The Patched domain protein PTR-4 is required for proper organization of the precuticular apical extracellular matrix. Genetics. 2021;219(3). doi: ARTN iyab132 10.1093/genetics/iyab132. PubMed PMID: WOS:000728164100006.

33. Serra ND, Darwin CB, Sundaram MV. Caenorhabditis elegans Hedgehog-related proteins are tissue- and substructure-specific components of the cuticle and precuticle. Genetics. 2024;227(4). doi: 10.1093/genetics/iyae081. PubMed PMID: WOS:001313022500001.

34. Wei H, Weaver YM, Yang CD, Zhang Y, Hu GL, Karner CM, Sieber M, DeBerardinis RJ, Weaver BP. Proteolytic activation of fatty acid synthase signals pan-stress resolution. Nat Metab. 2024;6(1):113-+. doi: 10.1038/s42255-023-00939-z. PubMed PMID: WOS:001135013100004.

35. Linkert M, Rueden CT, Allan C, Burel JM, Moore W, Patterson A, Loranger B, Moore J, Neves C, MacDonald D, Tarkowska A, Sticco C, Hill E, Rossner M, Eliceiri KW, Swedlow JR. Metadata matters: access to image data in the real world. J Cell Biol. 2010;189(5):777–82. doi: 10.1083/jcb.201004104. PubMed PMID: WOS:000278177500003.

36. Schindelin J, Arganda-Carreras I, Frise E, Kaynig V, Longair M, Pietzsch T, Preibisch S, Rueden C, Saalfeld S, Schmid B, Tinevez JY, White DJ, Hartenstein V, Eliceiri K, Tomancak P, Cardona A. Fiji: an open-source platform for biological-image analysis. Nature Methods. 2012;9(7):676–82. doi: 10.1038/Nmeth.2019. PubMed PMID: WOS:000305942200021.

37. Bai X, Smith HE, Golden A. Identification of genetic suppressors for a BSCL2 lipodystrophy pathogenic variant in Caenorhabditis elegans. Dis Model Mech. 2024. Epub 20240308. doi: 10.1242/dmm.050524. PubMed PMID: 38454882.

38. Bai X, Smith HE, Romero LO, Bell B, Vasquez V, Golden A. Mutation in F-actin Polymerization Factor Suppresses Distal Arthrogryposis Type 5 (DA5) PIEZO2 Pathogenic Variant in Caenorhabditis elegans. bioRxiv. 2023. Epub 20230724. doi: 10.1101/2023.07.24.550416. PubMed PMID: 37546771; PMCID: PMC10402071.

39. Bai X, Smith HE, Golden A. Identification of genetic suppressors for a BSCL2 lipodystrophy pathogenic variant in Caenorhabditis elegans. Dis Model Mech. 2024;17(6). Epub 20240416. doi: 10.1242/dmm.050524. PubMed PMID: 38454882; PMCID: PMC11051982.

40. Mok CA, Au V, Thompson OA, Edgley ML, Gevirtzman L, Yochem J, Lowry J, Memar N, Wallenfang MR, Rasoloson D, Bowerman B, Schnabel R, Seydoux G, Moerman DG, Waterston RH. MIP-MAP: High-Throughput Mapping of Caenorhabditis elegans Temperature-Sensitive Mutants via Molecular Inversion Probes. Genetics. 2017;207(2):447–63. Epub 2017/08/23. doi: 10.1534/genetics.117.300179. PubMed PMID: 28827289; PMCID: PMC5629315.

41. Hendi A, Mizumoto K. GFPnovo2, a brighter GFP variant for in vivo labeling in C. elegans. MicroPubl Biol. 2018;2018. Epub 20180906. doi: 10.17912/49YB-7K39. PubMed PMID: 32550394; PMCID: PMC7282520.

